# Individual variation of the dark-background-contingent upshift of gaze: effect of past habituation

**DOI:** 10.1101/628610

**Authors:** Oleg Spivak, Peter Thier, Shabtai Barash

**Author notes:** **Corresponding author:** Shabtai Barash, Dept. Neurobiology, Weizmann Inst. of Science, Rehovot 7610001, Israel.

## Abstract

We studied the shift of gaze direction induced by dark background in monkeys. Faced with large inter-individual variability, we asked how common the upshift is, and how the upshift size is distributed. Furthermore, we sought to reckon processes influencing the variability. Approaching these questions necessitates a large sample. Here we report data from 10 rhesus monkeys recorded in Tübingen, together with reported data from 4 cynomolgus monkeys studied in Rehovot. In all 14 monkeys, dark background induced upshift – but no systematic horizontal shift. The upshift might be thought of as a simple sensorimotor response; nevertheless, surprisingly, the monkeys’ previous experience appeared to have a decisive role in influencing the upshift’s size. All the monkeys were previously trained in tasks that involved vision and eye movements; by their previous training, the monkeys were naturally divided into two groups. Monkeys of the first, ‘bright-habituated’ group, previously trained in photopic, bright ambient-light conditions; monkeys of the second, ‘dark-habituated’ group previously trained mostly with isolated dots of light appearing in dim ambient lighting or in full darkness. The dark-habituated monkeys had a larger upshift than the bright-habituated: the groups were separated by a border-value such that 6/7 of the dark-habituated monkeys had upshift larger than the border, and 5/7 of the bright-habituated monkeys had upshift smaller than the border. Thus, the size of the dark-background-induced upshift largely reflects the extent to which a monkey is habituated to work in the dark. Though the upshift is reflex-like sensorimotor behavior, its amplitude largely reflects cumulative experience.

## Introduction

This study deals with the ‘upshift’, a puzzling behavioral phenomenon previously described in monkeys. As is well known, in bright light, if a single target appears on a computer monitor, monkeys facing the monitor direct their foveal line of gaze towards the target; the target’s image falls on the fovea. The behavior at issue in this study takes place after the bright ambient light is shut off, and the target’s visual background on the computer monitor is made dark. In this condition, at least some monkeys appear to look above the target rather than at it; more precisely, the foveal line of gaze is directed towards a location slightly above the target (Snodderly, 1987; Barash et al., 1998; Spivak et al., 2014). This is the dark-background-contingent upshift of gaze; for short, in this manuscript, ‘upshift’. The upshift is puzzling because it constitutes an exception to the rule that saccades and visual fixation movements always direct the target’s image to the fovea. What is beyond this exception?

### Is the upshift related to scotopic vision?

What is the function of the upshift? Is it but an idiosyncrasy of macaque sensorimotor processing, devoid of function? Because the upshift is observed when everything around the monkey (except the small target) is in darkness, it is tempting to speculate that the upshift is related to scotopic vision (Barash et al., 1998). Indeed, anatomical discoveries made around 1990 could be relevant to understanding the upshift. The dorsal retina was found to be rod-dense; rod density peaks at a retinal locus dorsal to the fovea, called rod hotspot (Packer et al., 1989) or dorsal rod peak (Wikler and Rakic, 1990; Wikler et al., 1990). Could the upshift be associated with the geometry of rod density? Perhaps scotopic vision could direct the target’s image to the rod peak rather than to the fovea, and that would reflect as an upshift of the foveal line of gaze (Barash et al., 1998).

However, most of the data regarding the upshift suggest a more complex relationship between behavior and anatomy. Exploring whether the upshift indeed relates to scotopic vision is possible only when vision is definitely scotopic, requiring a full interval of dark adaptation prior to testing. However, the monkeys previously tested were, by and large, not properly dark-adapted: often, testing the monkeys in the dark started shortly after being tested in bright illumination. Therefore, the testing transpired largely during the early phase of dark adaptation, when cones still dominate vision because rods are still saturated. Indeed, at least some monkeys react to an instantaneous offset of bright background illumination with an eye movement that shifts the eye to an upshifted fixation position. This reaction takes place briefly after background offset, with a latency similar to that of a saccade (see Figs 7,8 of (Barash et al., 1998)). At least in these monkeys, upshift appears to emerge almost as a reflex sensorimotor response to the change in the lighting of the animal’s environment, not unlike the almost-reflexive manner that saccades follow in photopic vision a target’s jump.

Thus, whether or not the upshift is present in scotopic vision, it is definitely not confined to scotopic vision. At least in some monkeys, it emerges clearly during the transition phase that briefly follows dark-background onset.

### Is the upshift a peculiarity of only some monkeys?

It had been obvious in previous studies, as in the new data presented here, that the upshift is a highly variable behavior pattern. In some monkeys, the upshift is large and salient, with the target falling well outside the fovea. However, in other monkeys, the upshift is smaller, and in some monkeys, it is not obvious if an upshift is present at all. Moreover, one study cast doubt about the very existence of the upshift (Goffart et al., 2006). These observations beg the question of reckoning the distribution of the upshift in the population. We translate the qualitative question of what fraction of the monkeys show an upshift to quantitative measures. What fraction of the population have a down-shift? A down-shift would be a dark-background mean fixation position lower than that with bright background. How does the up/down shift compare to left/right? Is there a systematic bias on both axis? On neither?

### The size of the upshift

As already mentioned, the upshift is highly variable between monkeys. We can measure the size of the upshift in degrees, reflecting the eye’s rotation. However, a pertinent consideration relates to the gaze-direction variability under bright ambient light, in photopic vision. Both intra-trial and inter-trial variability are present in photopic fixations. For evaluating the physiological significance of the upshift, the statistics of fixations in dark background makes sense only in reference to the variability of fixation in photopic vision. Towards this aim, we have defined an alternative measure for the size of the upshift, using an index calculated relative to the variability of fixation in photopic vision. Both absolute (degrees of rotation) and relative (index) measures of the upshift gave essentially the same results. Nonetheless, this similarity of results only begged the question: why is there such a large variability in the size of the upshift between individual monkeys? Perhaps the most intriguing result of the present study identifies a factor that appears to contribute heavily to this inter-individual variability of the upshift. Surprisingly, if we think of the upshift as a reflex-like behavior, this factor alludes not to the immediate past but to the experiences that the individual monkeys had during months preceding the testing for the upshift.

### Individual differences in monkey behavior

Addressing the questions of the present study requires a relatively large sample of monkeys. We have tested 10 monkeys for upshift in the recent years in Tübingen; we add the results of 4 previous monkeys tested in Rehovot (Barash et al., 1998). Altogether, we have a sample of 14 monkeys, comprising 2 species. This sample turned out to be large enough to yield answers to the questions we asked. Thus, a large sample allowed us to gain insights that could not be obtained with a standard sample of 2-3 monkeys.

N=14 is very large nowadays for controlled laboratory studies of monkey behavior, due to the external constraints. We have therefore tried in the discussion to come up with some thoughts regarding possible relevance of the present study for the common general situation, of having only 2 or 3 monkeys.

## Materials and methods

### Database

In Tübingen, we used 10 rhesus monkeys in this study (specified as T1 to T10). The monkeys had been previously trained in various oculomotor tasks. Some monkeys were used in electrophysiological recordings unrelated to this project. The other 4 *Macaca fascicularis* monkeys (R1 to R4) were previously studied in Rehovot. The results were published; here we used data included in this paper (Barash et al., 1998). All monkeys were male.

The procedures for the Rehovot monkeys were published (Barash et al., 1998). Therefore, here we will describe only the procedures related to the Tübingen monkeys.

### Animal preparation

All experimental procedures are standard and have been described in detail (Dash et al., 2012; Caggiano et al., 2013; Spivak et al., 2014). In brief, gaze direction was recorded using the scleral search coil method (Fuchs and Robinson, 1966; Judge et al., 1980). The monkeys were prepared for neurophysiological recordings (not part of the present study); their heads were painlessly immobilized by titanium head posts. Surgeries were performed under intubation anesthesia with isoflurane and nitrous oxide, supplemented by continuous infusion of remifentanil (1–2.5 (micro)g/kg1·h1) with full tight control of all relevant vital parameters. All procedures conformed to the National Institutes of Health Guide for Care and Use of Laboratory Animals and were approved by the local ethical committee (Regierungspräsidium Tübingen).

### Experimental setup

The experiments were conducted in a lightproof electrophysiological setup. The monkeys were seated in front of a cathode ray tube (CRT) screen at the distance of 30-40 cm from the screen center. The CRT was an Eizo Flexscan F730, 50-cm diagonal, displaying 1024×768 pixels at a frame rate of 60 Hz. The only source of light during the experiment was the monitor the monkeys were facing. Gaze direction (eye position) was sampled at 1000 samples per second. Data collection was performed using the open source measurement system *nrec*, https://nrec.neurologie.uni-tuT3ngen.de/nrec/nrec, created by F. Bunjes, J. Gukelberger and others. *nrec* is the standard recording system for behavior and electrophysiology at the Thier lab at Tübingen. The *nrec* output files were converted to *Matlab* (https://www.mathworks.com/, MathWorks, Natick, Massachusetts, USA). Post hoc data analysis was carried out in *Matlab*.

### Paradigm and training

A trial began with the appearance of small circular target spot at an unpredicted position on a screen. The size of the target was 0.02-0.1 deg. The monkeys had 1 s to make a saccade to the invisible fixation window centered on the target and to maintain the eyes within the fixation window for 1.5-2 s. The size of the window was small (2-3 deg radius) in bright-background trials and larger (5-15 deg radius) in dark-background trials – so that shifted fixation would not be interpreted by the computer as a fixation failure. The invisible fixation window was symmetric around the scotopic target, so that any shift, upwards or not, would have the same opportunity to be detected. On error the trial was aborted, and a new trial started; on correct performance the monkey was rewarded with a drop of water. It is noteworthy that the monkeys were rewarded for performing the task, but not for showing or not showing upshift. The upshift we report is spontaneous behavior, emerging in the conditions of the task.

### Visual stimuli

Target locations were arranged on 3 concentric circles of 5, 10 and 15 deg radii. The location of every target was selected in a randomly interleaved order, 1 target per trial. The experimental session consisted of a series of alternating blocks of dark-background and bright-background trials, usually 120 trials per block. Every experimental session began with a standard gaze direction calibration using bright background. The luminosity of the target was set to 70 cd/m^2^ but because in most of the cases it was very small (0.02 deg) it only spanned a few pixels.

Bright background was set at 7 cd/m^2^; dark background was set at such a level that for humans sitting in the experimental chamber for up to 3 hours, the dark-background monitor remained completely undiscernible.

### Analysis

To calculate the mean gaze direction during visual fixation that will represent each trial (Fig 1) we averaged across the interval of the last second of the trial. Although trial duration was variable, by taking into account the last second of fixation we made sure that at least 0.5 s elapsed from the time the eye entered the fixation window till the start of the time-interval over which gaze direction samples were taken for the calculation. Hence, during the 1 s interval used to calculate the trial’s representative fixation position all correction saccades were already completed, and the eyes were stably fixating. Trials with atypical gaze direction (less than 0.5%) were excluded from the analysis. For every monkey and every target location, we subtracted the mean bright-background gaze direction, mean of all bright-background trials, from each trial. Thus, the mean bright-background gaze direction is equivalent for an improved calibration of the direction of gaze while fixating the pertinent target. The shift of the dark-background trials from the mean bright-background gaze direction reflects the effect of the background illumination on gaze direction (Fig 1).

**Figure 1.**
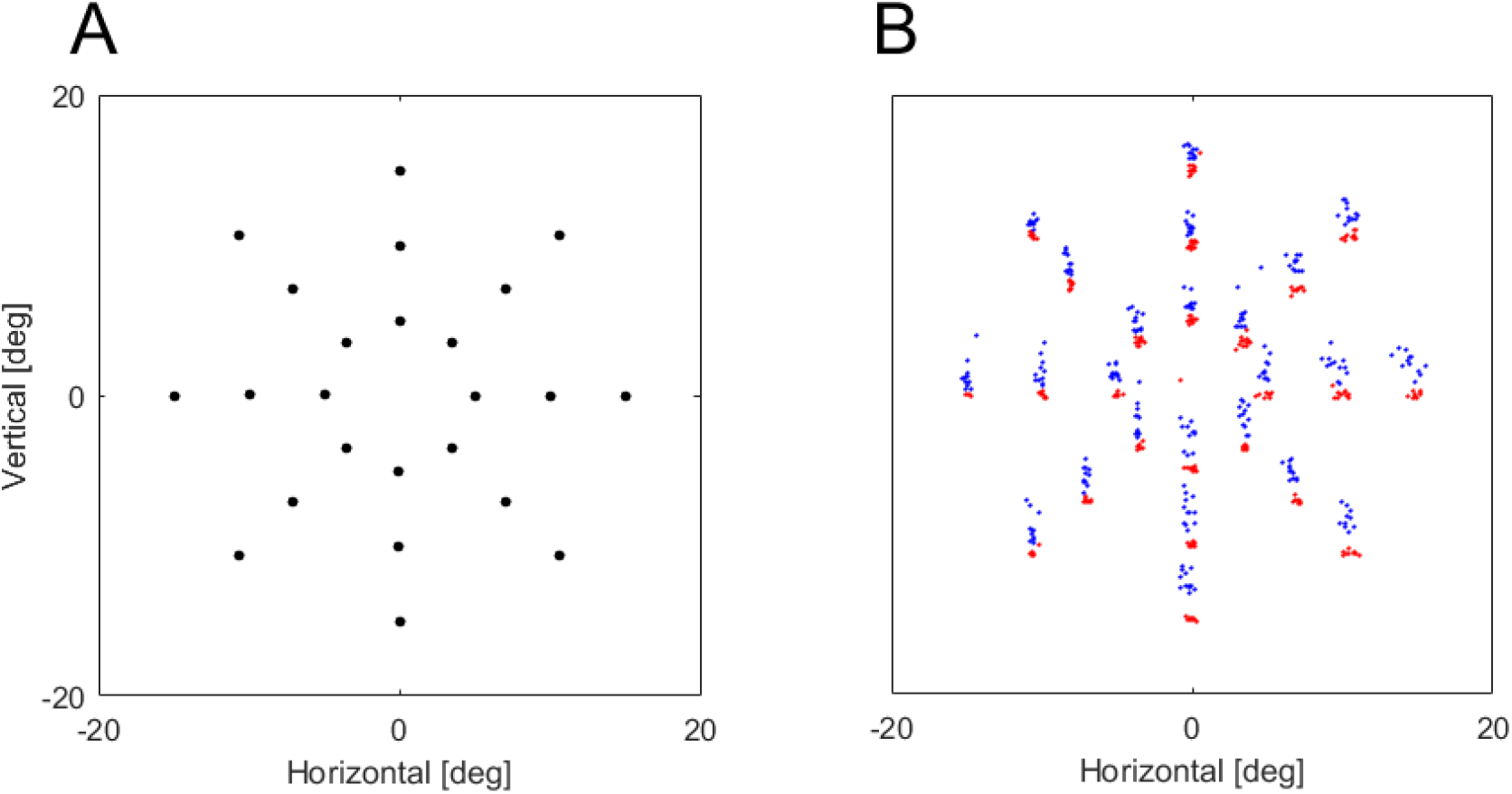
Target locations and illustration of the results from an example section. A) 24 possible target locations on the screen. The locations are arranged on 3 concentric circles with 5, 10 and 15 deg radii. (In the Rehovot monkeys the outer circle radius was 20 deg). B) Gaze direction data of one session of one monkey. Each dot represents the mean gaze direction during target fixation in one trial. Red dots stand for the bright-background trials and blue dots for dark-background trials. The locations of the red dot clusters closely correspond to the locations of the targets. Blue clusters are shifted upwards with respect to the red clusters illustrating the dark-background-contingent upshift. Note that in both conditions, the same targets were fixated. The only difference between the conditions was the background illumination.

## Results

We will begin by describing the procedures for acquiring the data, and for the analysis of each individual monkey’s data. Then we will move on to examine the results of the 14 monkeys together. The experimental procedure was previously described (Barash et al., 1998; Spivak et al., 2014); the analysis at the single-monkey level is a variation of that previously described (Barash et al., 1998; Spivak et al., 2014).

### Procedure, an example session, and definition of upshift

We studied visual fixation of small target spots; see Methods for details. There were two conditions: (1) bright background, and (2) dark background. The target spot was the same in both conditions – bright but very small, only a few pixels across. The stationary target spot was the only stimulus present on the screen. The background of the spot was uniform, gray (‘bright’) or black (‘dark’). Other than the computer screen in front of the subject, there were no light-sources in the experimental room and the room was light-tight. Thus, in the dark-background condition only the target was visible. We did not systematically record the state of dark adaptation during most of these recordings. Nonetheless, recordings in the dark-background condition typically took place shortly after the computer changed the screen’s background from bright to dark. At this time, vision is thought to be dominated by cones, with rods still largely saturated. The time-course of dark adaptation is in its initial, cone-dominated phase (Hecht et al., 1937; Reuter, 2011). Over the series of trials making up a block, targets appeared in 24 positions, which were arranged in 3 concentric circles of radii 5^0^, 10^0^, and 15^0^, illustrated in Fig 1A. (In the Rehovot monkeys, we used an outer circle of 20^0^, enabled by a slightly different display system). In Tübingen, a block typically consisted of 120 trials, with each target appearing in 5 trials, and the order of targets randomized. The Tübingen monkeys were previously trained in visual and oculomotor tasks; they performed the fixation task of the present study with little additional training.

For each trial, we calculated a mean fixation position by averaging each of the horizontal and vertical gaze direction components over the 1000 samples taken during the last 1s of fixation in that trial. Each dot in Fig 1B stands for the mean gaze direction of one trial. The red dots reflect fixation position in trials with bright background. There are 24 tight red clusters in Fig 1B and their configuration precisely reflects the configuration of the target positions in Fig 1A. Each cluster of red dots in Fig 1B reflects the set of trials sharing the corresponding target position (Fig 1B). Although the mean gaze directions vary a little from trial to trial, this variability is small, allowing the sets of trials sharing the same targets to form well isolated clusters of red dots in Fig 1B.

The blue dots of Fig 1B reflect the mean gaze directions in trials with dark background. The blue dots also partition into 24 clusters; each blue cluster is located slightly above a red cluster. The blue dots and the red dots in adjacent clusters reflect the mean gaze direction in trials with the same target. That the blue dots are above the red dots is the graphical expression of the upshift.

Thus, Fig 1B is marked by pairs of cluster, a red-dot cluster and a blue-dot cluster in each pair, with each cluster-pair corresponding to a target location. At issue is the deviation of gaze direction observed with dark background; that is, the shift of the blue cluster from the red, in each pair. We thus seek to define an gaze direction ‘shift’ on going from bright background to dark background. Towards that aim, for the trials of each target we define a base position as the mean gaze direction of all the bright-background trials directed to the target under consideration. Graphically, the base position is the center of mass of the target’s red cluster. For each trial with that target, that is, for each dot in the relevant pair of red and blue clusters, we define the trial’s shift as the vector that begins in the base position and ends at the dot corresponding to that trial. Here ‘shift’ refers to the trial’s deviation from the typical gaze direction while fixating the target at issue – that is, the deviation from the base position, as defined above. By studying shifts we can collect together the data from all targets together.

### Dark-background-contingent gaze shift for each of the 10 Tübingen monkeys

Fig 2 shows analysis of the dark-background-contingent shifts, separately for each Tübingen monkey. The analysis comprises a scatterplot of the trials’ shifts, and corresponding horizontal and vertical histograms. Table 1 lists the numbers of sessions/trials for each monkey. The histograms of the bright-background trial shifts are plotted in red and yellow, the dark-background trial shifts in blue and yellow. Yellow reflects the overlap of the two histograms. A red dot in a scatterplot reflects the mean gaze shift of a bright-background trial; a blue dot, the mean gaze shift of a dark-background trial. Graphically, the red dots occlude the blue; it is as if the blue dots are plotted first, and then the red dots in front of the blue.

**Figure 2.**
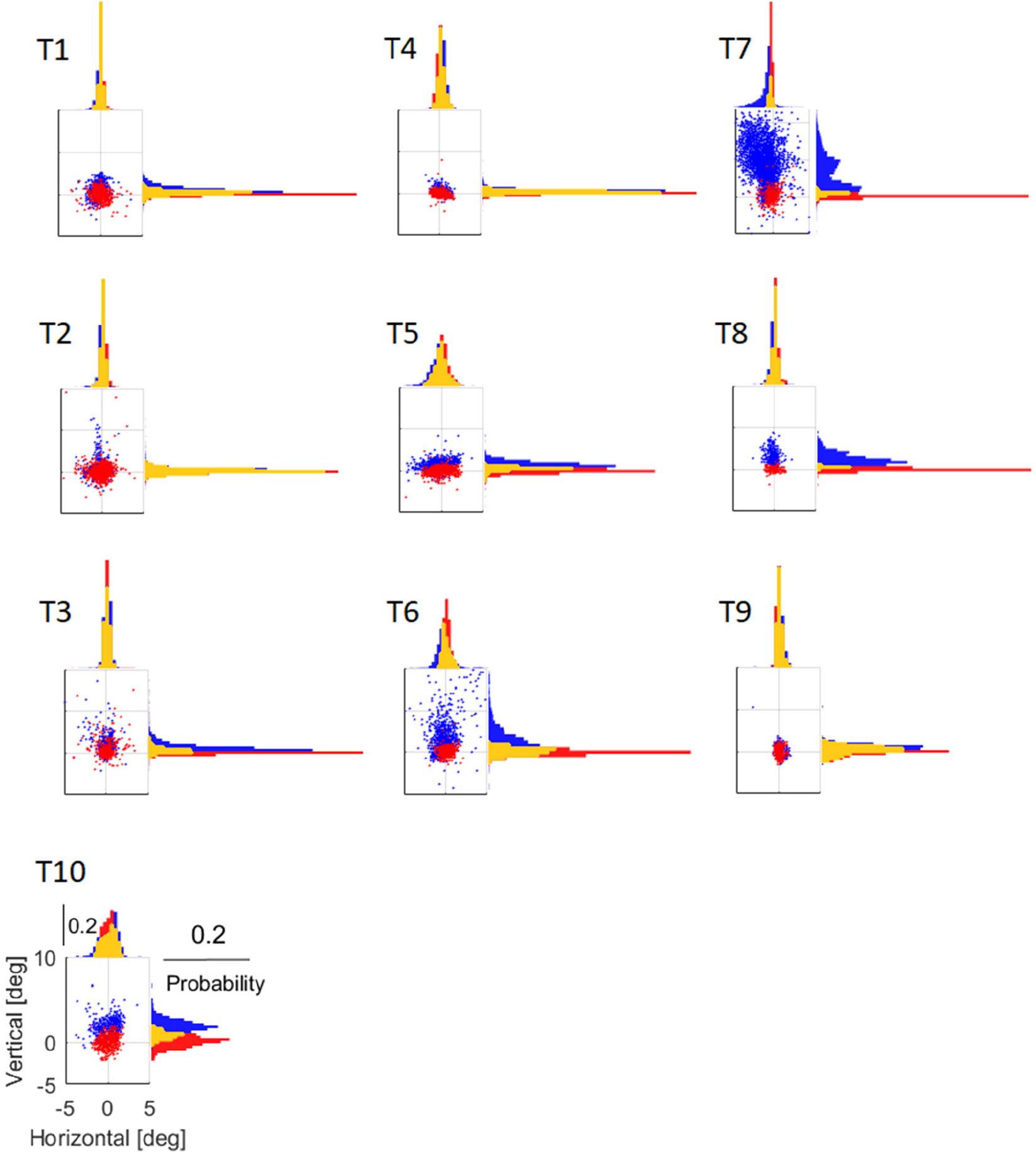
Gaze direction data with bright and dark backgrounds in the Tübingen monkeys. Mean gaze direction of the 10 Tübingen monkeys (T1-T10) during target fixation in dark and bright background trials, in all the sessions pulled together. Each data point on the scatterplots represent the difference between gaze direction in one trial and the approximated target location (see text). Red stands for the bright-background trials and blue for dark-background trials. Histograms of the vertical and horizontal components are displayed on the right and the top of each scatter plot respectively. Yellow bins depict the overlap between red and blue histograms.

**Table 1.**
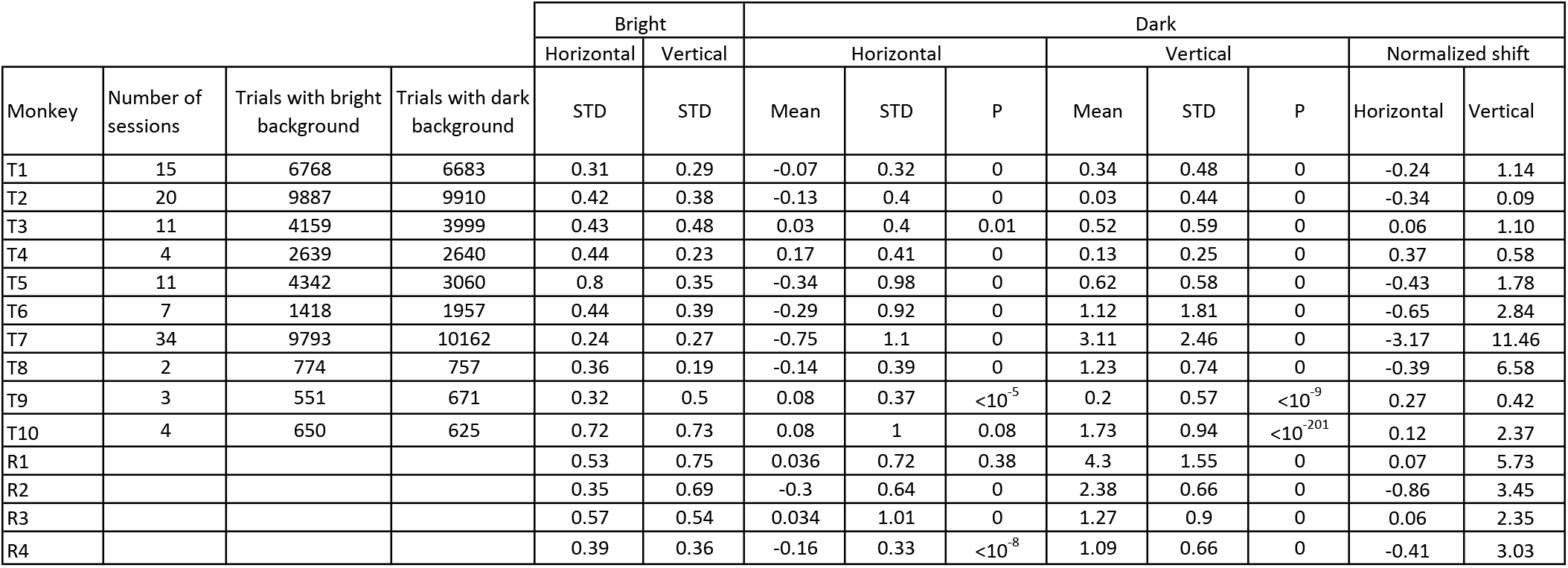

#### The bright-background shifts

For all of the monkeys, the red dots form a cluster. That the clusters are all centered on (0,0) is an outcome of our definition of a trial’s shift – the difference of gaze direction of the bright-background trials from their mean has itself a mean of zero. The histograms also show that both horizontal and vertical components of the bright-background shifts, that is, the red-yellow histograms, are centered on zero.

The shape and size of the red-dot clusters vary from monkey to monkey. In some monkeys the clusters are tight (for example, T8), whereas in others large (as in T2). In some monkeys the red clusters are elongated along the horizontal axis (as in Lennie), in others elongated along another orientation (as in T7), in yet others nearly circular (as in T1). These differences are likely to have multiple sources. Trial-by-trial variation of visual fixation position is very well documented (Cherici et al., 2012). Variability might stem also from other sources, including bringing together data from different target positions and data from different sessions (experimental days). Regardless of the causes for the shapes and sizes of the red clusters, in the present study are baseline. At issue here are primarily the blue dots – specifically, the way the blue dot clusters and blue-yellow histograms relate to the red and red-yellow.

#### The dark-background shifts

The blue dots in each of the scatterplots of Fig 2 form clusters (T7’s cluster is clipped so that the scales would fit the data of all the monkeys in Fig 2). Do the blue cluster look similar to the red? We make the following observations.

First, the blue clusters vary greatly from one monkey to another. In some monkeys the blue clusters are tight (for example, T9) whereas in others, much larger (T7).

Second, the overlap of the blue and red clusters also varies greatly from monkey to monkey. This is salient in the histograms. The overlap between the histograms of the vertical shifts is colored yellow. In some monkeys yellow predominated the vertical histograms, with blue and red barely showing (for example, T2). In other monkeys the yellow area is minor compared to the red and blue (T8).

Third, the horizontal extent of clusters also varies from monkey to monkey, but for most monkeys the yellow area in the horizontal histograms is large; the bright-background and dark-background mean fixation positions largely overlap each other.

Fourth, scanning the general shape of the blue clusters and comparing them to the red, the most prominent feature coming out is a shift of the blue clusters upwards from the red. The size of the shift varies: in some monkeys the blue dots are mostly above the red (T7), in others the shift is smaller. Importantly, there is no systematic analogous horizontal shift. In some monkeys there is a horizontal shift (for example, note how the blue cluster of T6 surrounds the red on the left). However, even in these shifts appear secondary to the vertical shifts. There is no monkey with a horizontal shift similar to the distinct vertical shift in Lennie, or even in T10.

### Normalized shifts

We seek to compare the distance between the blue and red data in different monkeys. The data in Fig 2 can be used for this comparison, but a careful look at the Figure brings up a pertinent question. Given the very different variability in bright-background fixations, that is, the very different size of the red clusters, are degrees of visual angle appropriate for measuring the distance between the blue and red clusters? One could argue that an absolute measure such as degrees is inappropriate because it is invariant of an individual monkey’s variability of the red dots. If two monkeys have an upshift of, say, 5 deg, but one monkey’s fixations in bright background are very tight (say, standard deviation 1 deg) whereas the other monkey’s fixations in bright background are loose (say, standard deviation 4 degrees), one might argue that the same absolute shift of 5 deg is physiologically significant in monkey 1 but less significant in monkey 2.

To counter such arguments we defined a normalized shift, in the following way. The normalization is computed separately for the horizontal and vertical dimensions, separately for each trial. The normalized shift is defined as a trial’s shift, in degrees visual angle, divided by the standard deviation of the red dots, the bright-background fixations. Because the unit of the standard deviation of the red dots is also degrees, the resulting normalized shift is dimensionless. A value of 1 for the normalized shift of a trial indicates that the trial’s gaze is shifted 1 standard deviation of the bright-background trials from the bright-background mean fixation location.

Fig 3 shows the same data as Fig 2 but in normalized shifts. Although qualitatively the same data, the normalized shift is more distinct. This holds for monkeys with small shifts (T9, note the blue vertical line beyond the yellow in the vertical histogram), and monkeys in which the shift was visible also in visual degrees (compare the histograms of T10 in Figs 2 and 3).

**Figure 3.**
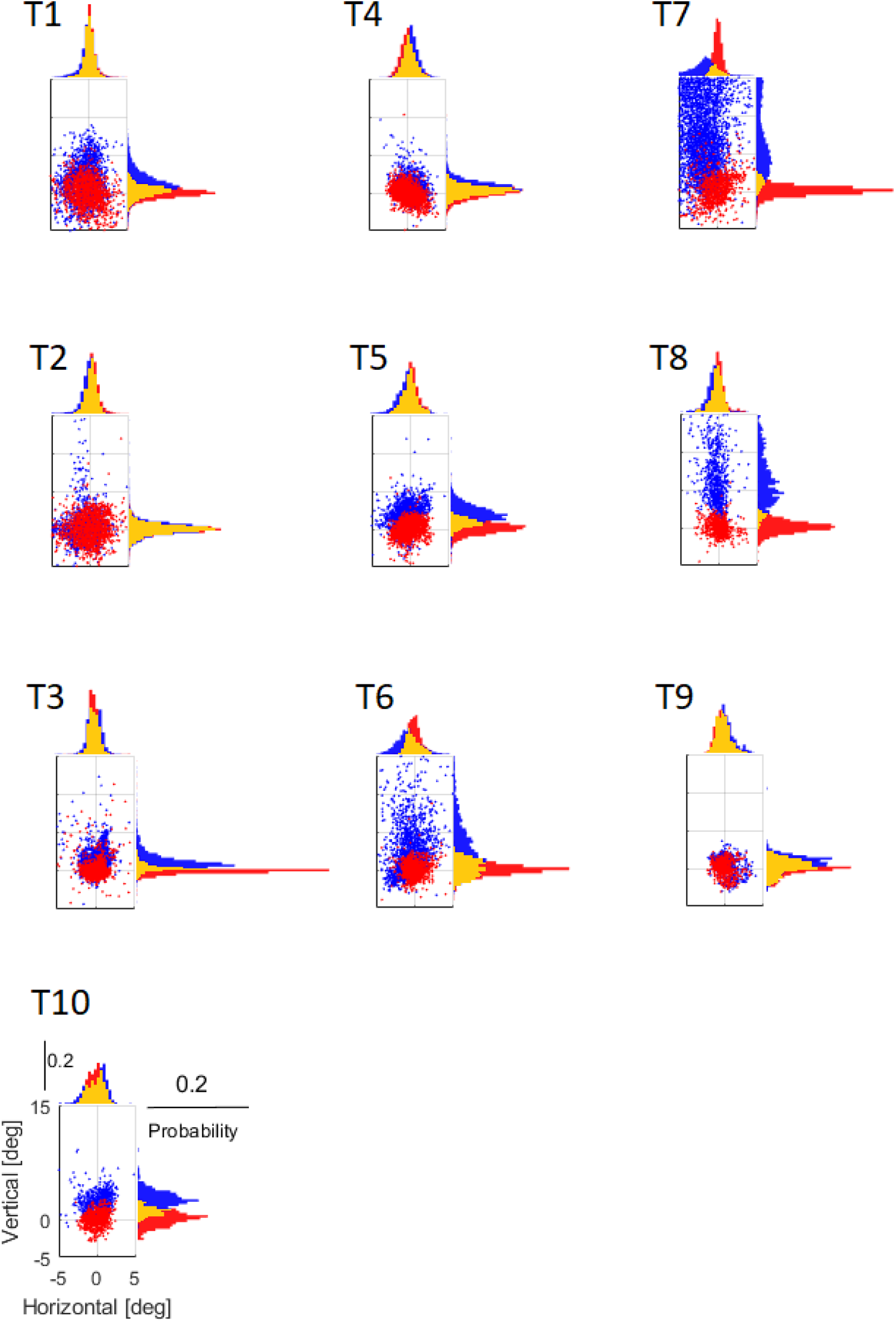
Normalized gaze direction in the Tübingen monkeys. Same data as in figure 2, normalized with respect to the variability of each monkey’s gaze direction in photopic vision. See text for details.

### Comparison of all monkeys

We now proceed to the study of the 14 monkeys together, the 10 Tübingen monkeys studied above and the 4 Rehovot monkeys from the 1998 paper. The data is listed in Table 1. The table lists for the bright background only the horizontal and vertical standard deviation; the mean is (0,0), by definition. For the dark data, both mean and standard deviation are included, for horizontal and vertical, as well as the normalized shift. The p-value relates to the difference of the mean from zero.

#### Distribution of the shifts

The first major result is a confirmation of the existence of dark-background-induced upshift. In fact, the commonality of upshift surprised us. In all 14 monkeys, the vertical shift is positive (that is, directed upwards). In contrast, there was no systematic horizontal shift. In 8 out of the 14 monkeys the horizontal shift was negative (leftwards); in the other 6 monkeys the horizontal shift was positive (rightwards). The values are listed in Table 1.

Fig 4 illustrates the distribution of the normalized horizontal and vertical shifts. The horizontal shift histogram is colored olive. Most of the horizontal shifts are close to zero: the horizontal shift of 13 monkeys is either on the bin centered on 0 or a single bin away from it.

**Figure 4.**
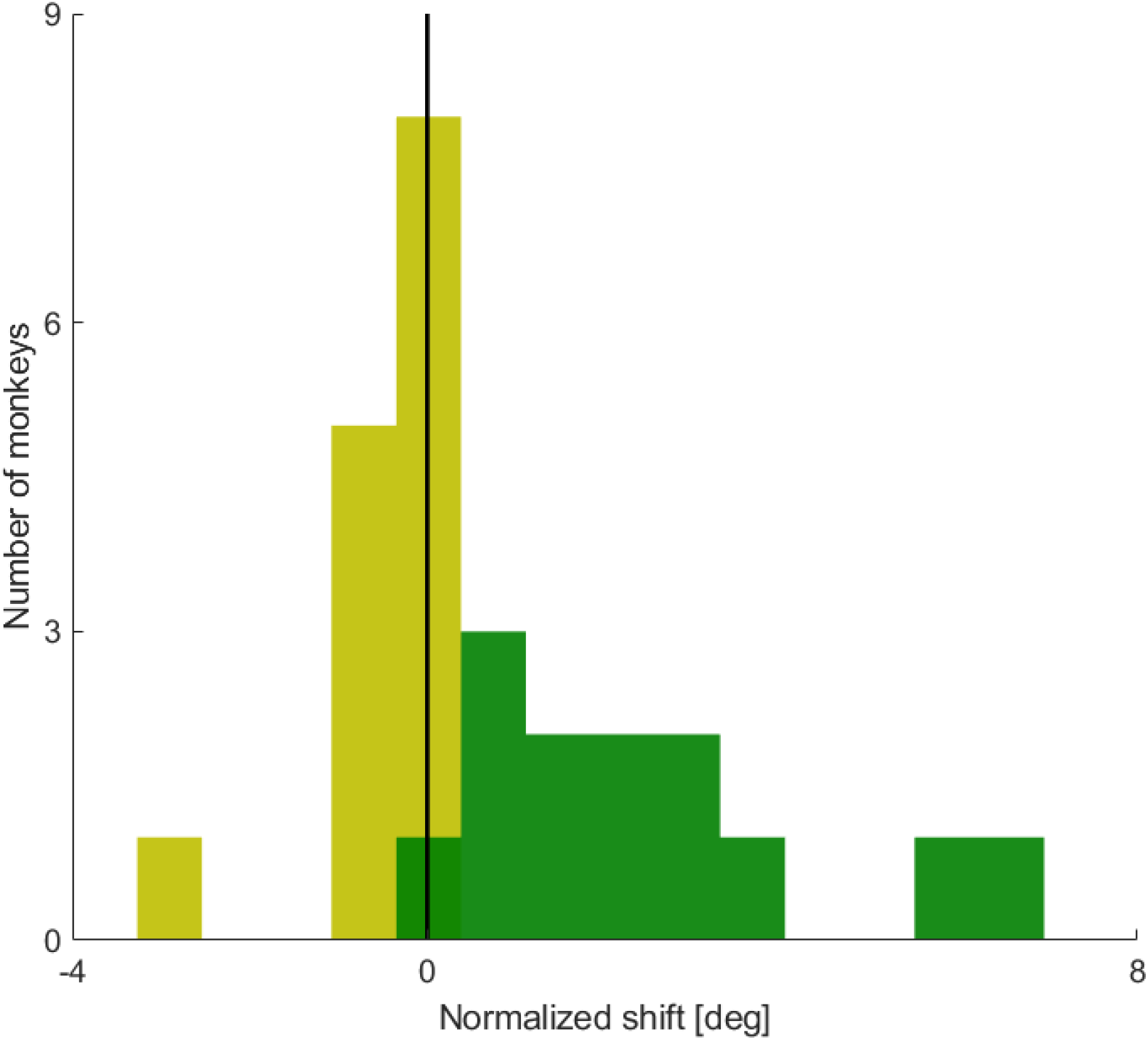
Horizontal and vertical shifts in the 14 monkeys. Histograms of the mean horizontal (olive) and vertical (green) normalized shift of the 14 monkeys. The black vertical line is at zero, signifying absence of shift. The horizontal shifts are close to zero; the vertical shifts are all positive, many of them large.

The dark green histogram reflects the distribution of the vertical shifts. A single monkey has a vertical shift in the bin centered on zero; thus, the mean vertical components of the dark- and bright-background fixations are very close, though the dark-background fixations are somewhat higher. Three more monkeys have a shift in the next bin. The other 10 monkeys’ vertical shifts fall in farther bins, reflecting their larger absolute values. Thus, the dark-green histogram shows that there is an upshift; its size varies from small in some monkeys to large in others. There is no horizontal analogue..

Figure 5 illustrates a scatterplot of the absolute (Fig 5A) and normalized (Fig 5B) shifts of all the monkeys. The Rehovot monkeys from the 1998 study listed as R1,…,R4 correspond to Monkey 1,…,Monkey 4 in that study. There are some differences between the scatterplots of the absolute and normalized shifts. For example, in the absolute plot (Fig 5A) R1 is shifted upwards of T7, but because of its R1’s inter-trial variability, in normalized shifts T7 has a greater upshift than R1.

**Figure 5.**
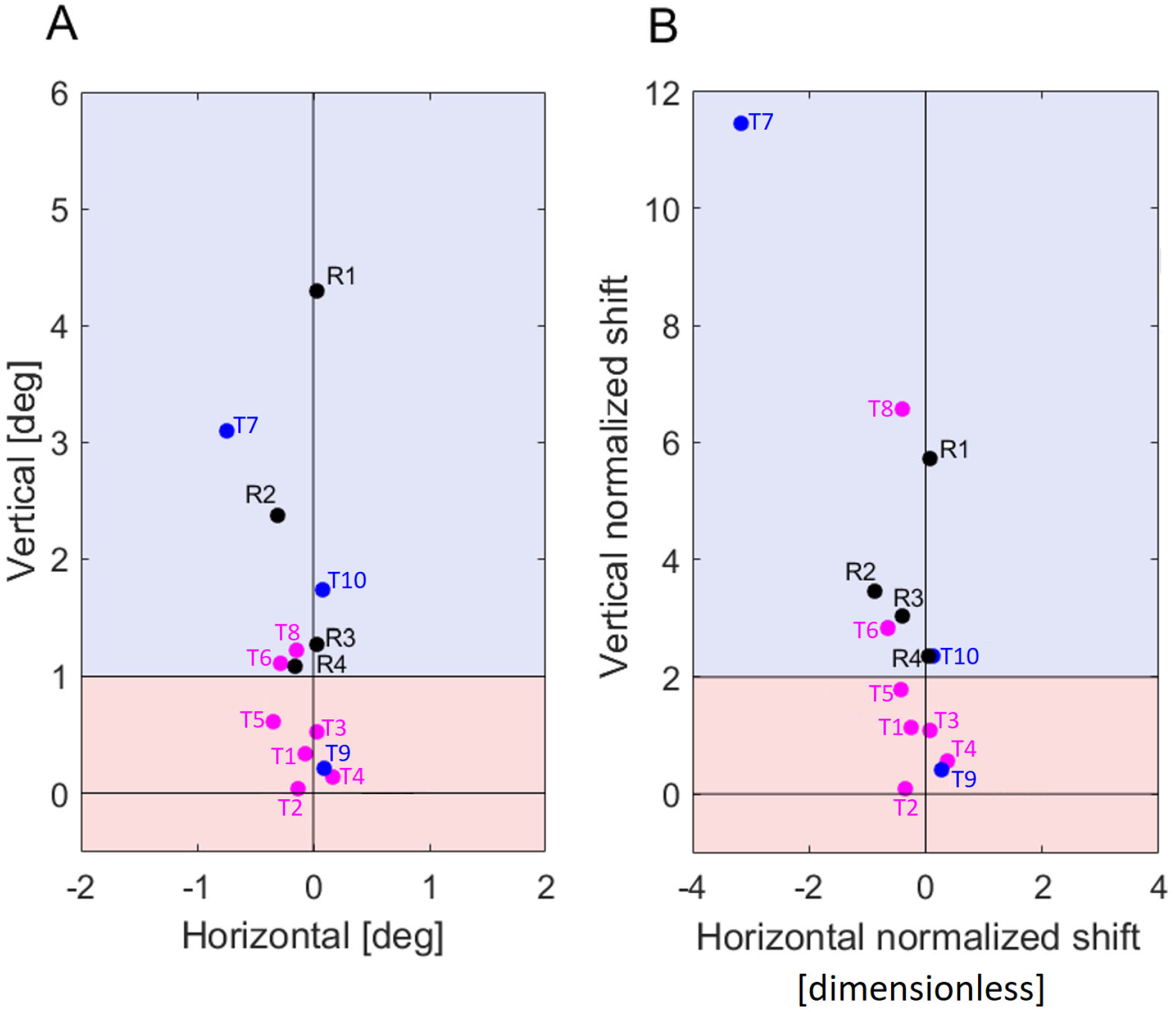
Upshift size largely reflects a monkey’s previous experience. A) Mean shifts of the 14 monkeys (dark-background mean gaze direction minus bright-background mean gaze direction). Each monkey’s ID is marked next to its dot. T1-T10 depict Tubingen monkeys, R1-R4 depict Rehovot monkeys. The bright-habituated monkeys are illustrated with pink color. The Tübingen dark-habituated monkeys are illustrated with blue color. The 4 dark-habituated Rehovot monkeys are illustrated with black color. Low upshift range (< 1 deg) is shown with reddish background, high upshift range (> 1deg) with bluish. Note that most of the dark-habituated monkeys are in the high-upshift range, whereas most of the bright-habituated monkeys are in the low-upshift range. The black vertical lines mark the horizontal and vertical meridians, and the division between low and high upshift ranges. B) Same data as in A, but normalized. Note that the division between the low and high upshift ranges is located at 2, reflecting 2 times standard deviation of each monkey’s gaze direction in photopic vision.

Two points predominate the scatter plots. First, in both scatterplots the monkeys are arranged on a rather narrow strip surrounding the vertical meridian. Second, in both scatterplots the monkeys are all represented in the half-plane above the horizontal meridian. These two points together show at the level of a large sample that the dark-background-induced upshift is ubiquitous and robust in macaque monkeys.

Another point that the scatterplots of Fig 5 show is that there is a lot of individual variability in upshift size. Is there any explanation for this variability?

#### Grouping based on experience in darkness

The Tübingen monkeys that we tested could be divided to 2 groups, according to the studies they had taken part in before we tested them with the upshift. Some of the studies were conducted mostly in the dark; for example, studies of cerebellar activity related to saccadic adaptation. Other studies were conducted with bright, daylight-intensity ambient light; for example, studies related to the perception of others’ gaze direction. We call these groups dark-habituated and bright-habituated, respectively. In Fig 5, the bright-habituated monkeys are listed in pink; the dark-habituated in blue. The Rehovot monkeys are illustrated in black. The Rehovot monkeys were all dark-habituated, trained in tasks such as memory-guided saccades with dark background. Altogether, we have 10 *Macaca mulatta* monkeys from Tübingen, 4 *Macaca fascicularis* from Rehovot, adding up to 14 monkeys. Out of these, 7 monkeys were bright-habituated (T2, T4, T1, T3, T5, T6, T8), and the other 7 dark-habituated (T9, T10, T7, R1, R2, R3, R4).

When we look at Fig 5 we see that in both absolute and normalized upshifts, the dots in the scatterplots can be divided into two ranges by setting a horizontal line as boundary between the ranges. In the absolute shifts (Fig 5A) we set this border line at 1 deg, and in the normalized shifts at 2. We call these two ranges low-upshift and high-upshift, correspondingly. In the low-upshift range there are 6 monkeys. Five of these are bright-habituated. In the high-upshift range there are 8 monkeys. Six of them are dark-habituated. Conversely, of the 7 bright-habituated monkeys, 5 (71%) are in the low-upshift range. Thus their upshift is less than 1 deg in size and less than twice the standard deviation of their vertical bright-background fixations. Of the 7 dark-habituated monkeys, 6 (86%) are in the high-upshift range; their upshift is greater than 1 deg, and greater than two times the standard deviation of the vertical component of their bright-background fixations.

## Discussion

In this study we revisit the upshift of gaze-direction, which is induced in monkeys by dark background. At issue is gaze direction of monkeys looking at small bright targets. The notion ‘upshift’ refers to the change of gaze-direction that reflects target luminosity; ‘shift’ is the gaze direction in dark background compared to bright background, and ‘upshift’ relates to the observations made in previous studies, using few monkeys, that the shift is primarily in the upward direction. In other words, the observation was that the eyes of the fixating monkeys are directed at the targets when the background is bright, but above the target when the background is dark.

### The upshift is a spontaneous, automatic behavior

The upshift is spontaneous behavior. The monkeys were never rewarded for shifting their gaze, neither upwards, nor in any other direction. If the background changes from bright to dark at once, the upshift follows briefly; at least in some monkeys the eyes then move upward in saccade-like movements, with brief reaction time, to the upshifted position (Barash et al., 1998). If the background then turns bright again, the upshift is abolished, typically with a similar saccade-like step, directed downwards. When we first came upon the upshift, we thought it was an artefact and tried to train it away. The only result was that the monkeys were quickly frustrated, and stopped working (Barash et al., 1998). Thus, the upshift is an automatic behavior that apparently is unrelated to the rewards the animals receive. The implicit, internal reward driving this behavior might be, as with many eye movements, the effect on the monkey’s visual performance.

### The upshift is ubiquitous

A commonly held tenet is that the core of visual fixation is foveation; in more detail, that the eyes are positioned during fixation so that the location of interest in the scene’s image (‘target’) falls on the fovea. The upshift is intriguing because it violates this foveation tenet. When a monkey has a small upshift, the target’s image only partially falls on the fovea. When a monkey has a moderate or large upshift, the target falls in the dorsal retina, away from the fovea. Thus, upshifted fixation does not foveate the target. Because the upshift is intriguing, the first question faced by the present study was – how general is the upshift? We computed the mean shift from all the monkey’s trials. We asked: what fraction of our sample of monkeys show positive vertical shift, that is, upshift? The answer – all the 14 monkeys we tested had upshift.

In all 14 monkeys, mean gaze was directed in dark-background fixations above its direction in bright-background fixations. This systematic bias upwards had no analogy in the horizontal component of gaze direction: in 8 monkeys the shift was leftwards, in 6 monkeys rightwards. Thus, the existence of the upshift is very robust. We observed it in 2 species, in laboratories far apart. (Although all monkeys were male, we expect the upshift to be gender-invariant). All in all, we were surprised by how ubiquitous the upshift is. This ubiquity emerges from our large sample. The large sample was necessary for obtaining this results.

### Upshift size is variable in both raw and normalized coordinates

When we consider the categorical yes/no question of whether the monkeys have upshift, we get the same answer in all the monkeys. However, if we instead consider the size of the mean upshift, the monkeys are highly variable. This variability is evident in numbers (mean dark over standard-deviation of bright column in Table 1). The variability is also evident by just looking at the scatterplots and histograms, in both of their versions, raw (Fig 3) and normalized (Fig 4). In photopic vision, monkeys as humans vary in the precision of their visual fixations (Cherici et al., 2012) This variability of gaze direction takes place when the background is bright, having nothing to do with the effect of dark background; therefore, it is useful to modulate the upshift, the difference of mean gaze directions in dark versus bright backgrounds, to a measure of the variability of a monkey’s photopic gaze direction. Hence the definition of the normalized shift. We separately normalized the horizontal and vertical components of the upshift. Study of normalized gaze confirmed the main results; indeed, the upshift is clearer after the normalization (compare Figs 2 and 3).

### The possible relationship of the upshift to scotopic vision is complex

Is the upshift related to scotopic vision? The answer is rather complex. On the one hand, the upshift appears to be associated with the retinal geometry of rod density. Rod density is high in dorsal retina. Dorsal to the fovea, rod density increases, peaking at a location called dorsal rod peak (DRP) (Wikler and Rakic, 1990; Wikler et al., 1990) or Rod Hotspot (Packer et al., 1989; Curcio et al., 1990). On the other hand, scotopic vision does not immediately follow an instantaneous offset of bright ambient illumination (such as the bright background in our study). Full dark adaptation takes as long as 45 min in the dark. The measurements reported here, in (Barash et al., 1998) and in other studies of the upshift were not subject to controlled dark adaptation and were mostly taken during the first phase of the dark-adaptation time-course, while vision is thought to be dominated by cones because the rods are still saturated. Moreover, at least in some monkeys, the shift of the eyes upwards briefly follows an instantaneous offset of the background; the upshift emerges in these conditions with a brief jump of the eyes, which might be a saccade (see Figs 7,8 of (Barash et al., 1998)). Thus, monkeys shift their gaze upwards within 1 s after the onset of dark background – long before the onset of scotopic vision. It transpires that upshift is a trait of the transition from photopic vision to dark-adjusted vision. Does this adjustment to darkness reflect an aspect of scotopic vision? This question goes beyond the scope of the present study; here we concentrate on the data we do have from 14 monkeys. This large sample makes it possible to ask questions that hard to approach with the small sample size common nowadays in monkey behavior studies.

### The dependence of upshift size on past experience

Perhaps the most important result of the present study is that much of the inter-individual variability of the upshift’s size can be explained by the monkeys’ experience prior to the period they were tested in the present study. There is a clear and significant difference between the two groups, one group encompassing the monkeys that were previously trained in photopic conditions and the other group encompassing monkeys that were previously trained with small target spots, mostly in the dark. Most of the monkeys in the bright-habituated group had mean upshift of less than 1 deg, and less than twice the standard deviation of the photopic fixations. Hence, although the fixations in the bright-habituated group differ significantly from photopic vision’s foveation movements, even when the background is dark their foveas partially overlap the projection of the target’s image on the retina. In contrast, in the dark-habituated monkeys, the target’s image barely overlaps the fovea or entirely falls out of the fovea. The clustering into the two groups is statistical; both groups have exceptions (T8 and T9, respectively) but the exceptions only substantiate the rule. To sum up, only by looking at the entire large sample do these inter-individual differences in upshift size emerge.

### Dependence on less recent past

Thus, upshift size crucially depends on the monkey’s past habits, with the formative period lasting months before we started to test monkeys for upshift, sometimes years. This dependence on a form of long-term memory is non-trivial. Well studied forms of motor learning, for example, saccadic adaptation, appear to depend primarily on the monkey’s recent performance. The upshift appears to be a stereotyped response that seems to be low-level. But if it were truly entirely low-level, we would expect the upshift to reflect the current sensory stimulation, or recent sensory stimulation (such as the state of light / dark adaptation). The dependence on much less recent past suggests that the upshift is not a pure bottom-up process but a process strongly modulated by top-down influences. Understanding this top-down modulation is out of the scope of the present study.

### What all that might suggest for 2- or 3-monkey experiments

Finally, cautiously, we would like to suggest that the results of this study might be relevant also for investigators of unrelated monkey behavior. It might be useful to bear in mind not only the monkeys’ performance, but also their previous training and life conditions. Perhaps in principle self-evident, this deliberation is not always practiced, and our results suggest that it might at times be more pertinent than initially assumed.

## Acknowledgements

To be added.

